# IL-23 signaling prevents ferroptosis-driven renal immunopathology during candidiasis

**DOI:** 10.1101/2021.12.19.473386

**Authors:** Nicolas Millet, Norma V. Solis, Diane Aguilar, Michail S. Lionakis, Robert T. Wheeler, Nicholas Jendzjowsky, Marc Swidergall

## Abstract

During infection the host relies on pattern-recognition receptors to sense invading fungal pathogens to launch immune defense mechanisms. While fungal recognition and immune effector responses are organ and cell type specific, during disseminated candidiasis myeloid cells exacerbate collateral tissue damage. However, the complex interplay between protective antifungal immunity and immunopathology remains incompletely understood. The β-glucan receptor ephrin type-A 2 receptor (EphA2) is required to initiate mucosal inflammatory responses during oral *Candida* infection. Here we report that Epha2 promotes renal immunopathology during disseminated candidiasis. EphA2 deficiency leads to reduced renal inflammation and injury. Comprehensive analyses reveal that EphA2 limits IL-23 secretion in dendritic cells, while IL-23 signaling prevents ferroptotic myeloid cell death during infection. Further, ferroptosis aggravates inflammation during infection, while at the same time reducing the fungal killing capacity of macrophages. Thus, we identify ferroptotic cell death as a critical pathway of *Candida-*mediated renal immunopathology that opens a new avenue to tackle *Candida* infection and inflammation.

## Introduction

The first step in mounting an antifungal immune response is the recognition of extracellular pathogen-associated molecular patterns (PAMPs) of invading organism, such as *Candida albicans*, by various families of soluble and membrane-bound pattern recognition receptors (PRRs) ^1,2^. Following recognition, the effective control of *C. albicans* relies on several effector mechanisms and cell types to ensure fungal clearance ^3-9^. In contrast to mucosal candidiasis, in which IL-17-producing lymphocytes are crucial for host defense ^10-14^, effective immunity during disseminated candidiasis relies on myeloid phagocytes ^2,15-17^. Although myeloid phagocytes are crucial for host defense during disseminated candidiasis, their functions that are aimed to control fungal infections may also come at the cost of immunopathology ^18,19^. In fact, excessive neutrophil accumulation in tissues late in the course of infection is deleterious in mouse models of disseminated candidiasis ^17,20^.

*C. albicans* is known to induce tissue injury and host cell death ^17,18,21^, e.g. in the kidney, a major target organ during disseminated candidiasis. Regulated host cell death (RCD) results in either lytic or non-lytic morphology, depending upon the signaling pathway ^22^. Apoptosis is a non-lytic, and typically immunologically silent form of cell death ^23^. On the other hand, lytic cell death is highly inflammatory ^23-26^, and includes necroptosis (alternative mode of RCD mimicking features of apoptosis and necrosis ^27^), pyroptosis (RCD driven by inflammasome activation ^22^), and ferroptosis (iron- and lipotoxicity-dependent form of RCD ^25^). Inflammatory RCD depends on the release of damage-associated molecular pattern (DAMPs) and inflammatory mediators ^28^. RCD is increasingly understood to benefit the host ^29^, and *C. albicans* is known to induce inflammatory RCDs, such as necroptosis, and pyroptosis to promote inflammation ^30,31^. Indeed, deficiencies in these pathways accelerate disease progression during fungal infection ^31,32^. However, excessive inflammation results in renal immunopathology during candidiasis suggesting that other mechanisms or RCDs fine-tune immunopathology and fungal control. Emerging data from various studies indicate an essential function of non-classical β-glucan recognition during fungal infections ^33-38^. We recently found that EphA2 acts as a β-glucan receptor in the oral cavity that triggers the production of pro-inflammatory mediators via STAT3 and MAPK on oral epithelial cells, while EphA2 induces priming of neutrophil p47^phox^ to increase intracellular reactive oxygen species (ROS) production to enhance killing of opsonized *C. albicans* yeast ^33,34^. Although the function of this novel β-glucan receptor EphA2 is well established during oral mucosal *C. albicans* infection ^33,34,39-41^, the role of EphA2 during disseminated candidiasis is unknown.

## Results

### EphA2 deficiency increases tolerance during disseminated candidiasis

Being the core of the immune response, professional immune cells act as the most effective weapon to clear invading fungi. Dectin-1/CLEC7A is a major PRR of the C-type lectin family, predominantly expressed on myeloid-derived cells. Classical β-glucan recognition by Dectin-1 activates fungal phagocytosis and the production of pro-inflammatory cytokines ^4,42^. Consistent with previous findings ^43,44^, Dectin-1 deficiency results in increased mortality in a mouse model of disseminated candidiasis (**Fig. 1A**). Although EphA2 recognizes β-glucan ^33,38^, and EphA2 deficiency results in increased susceptibility to oral fungal infection ^33,34,40^, *Epha2*^−/−^ mice were more resistant during lethal *C. albicans* challenge (**Fig. 1B, C**). Since EphA2 is expressed on both stromal and hematopoietic cells ^33,34,37,45,46^, we generated bone marrow (BM) chimeric mice and determined their resistance to disseminated candidiasis (**Fig. S1**). Both, *Epha2*^*+/+*^ mice reconstituted with *Epha2*^*–/–*^ BM (knockout (KO)→wild-type (WT)) and *Epha2*^*–/–*^ mice reconstituted with *Epha2*^*+/+*^ BM (WT→KO) were more resistant during HDC compared to *Epha2*^*+/+*^ mice reconstituted with *Epha2*^*+/+*^ BM (WT→WT) (**Fig. 1D**). However, these chimeric mice were more susceptible than *Epha2*^*–/–*^ mice reconstituted with *Epha2*^*–/–*^ BM (KO→KO), which recapitulated the phenotype observed in global *Epha2*^*–/–*^ mice (**Fig. 1B**), suggesting that EphA2 deficiency within cells of both, the hematopoietic and stromal compartments, is required for full protection against disseminated candidiasis.

**Figure 1.**
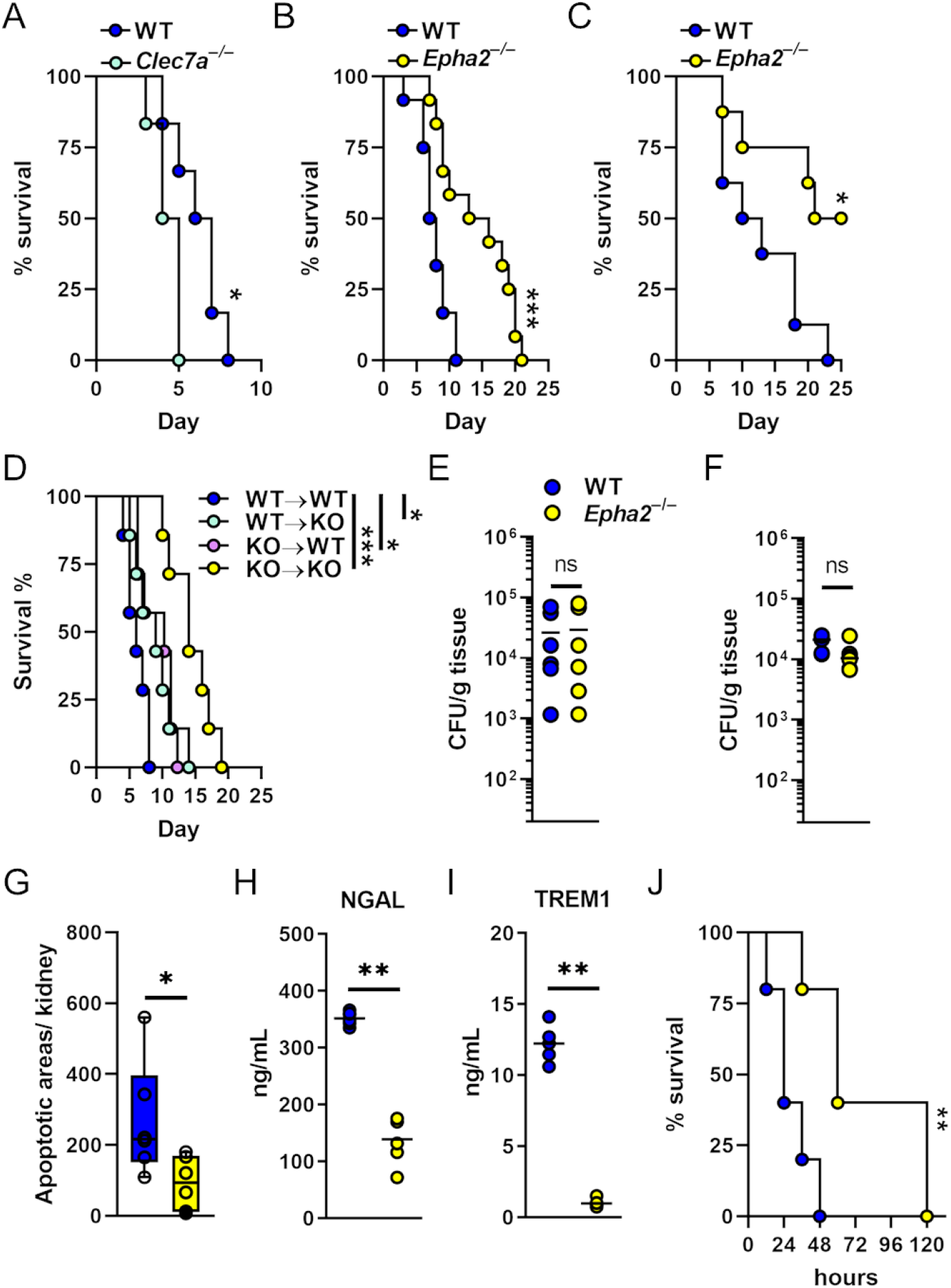
EphA2 promotes disease progression during disseminated candidiasis. **A** Survival of wild-type and *Dectin-1*^−/−^*(Clec7a*^−/−^) mice infected intravenously with 2.5×10^5^ SC5314 *C. albicans*. N=6; two independent experiments. **B**-**C** Survival of wild-type and *Epha2*^−/−^mice infected intravenously with 2.5×10^5^ (**B**; N=12; two independent experiments) or 1.25×10^5^ (**C**; N=8; two independent experiments) SC5314 *C. albicans*. **D** Survival of bone marrow chimeric mice following infection with 2.5×10^5^ SC5314 *C. albicans*. N=7. *Epha2*^*+/+*^ mice reconstituted with *Epha2*^*+/+*^ BM (WT→WT), *Epha2*^*+/+*^ mice reconstituted with *Epha2*^*–/–*^ BM (KO→WT), *Epha2*^*–/–*^ mice reconstituted with *Epha2*^*+/+*^ BM (WT→KO), and *Epha2*^*–/–*^ BM to and *Epha2*^*–/–*^ mice (KO→KO). Statistical significance is indicated by ^*^ *P<*0.01, and ^***^ *P<0,001*. Mantel-Cox Log-Rank test. Kidney fungal burden of infected mice after **E** 4 days and **F** 12 hours of infection with 2.5×10^5^. Results are median (N=6) of two independent experiments. ns, no significance. Mann-Whitney Test. **G** Apoptotic areas per kidney after 4 days of infection determined by TUNEL staining. N=6; two independent experiments. Statistical significance is indicated by ^*^*P<*0.05; Mann-Whitney Test. **H** Serum NGAL and **I** TREM1 levels after 3 days of infection. N=6; two independent experiments. ^**^*P<*0.01; Mann-Whitney Test. **J** Survival of wild-type and *Epha2*^*-/-*^ mice injected intraperitoneally with 750 mg/kg of zymosan. N=5; two independent experiments. Statistical significance is indicated by ^**^, *P <* 0.01. Mantel-Cox Log-Rank test

Kidneys are a primary target organ of *C. albicans*, and invasion into the kidney medulla leads to loss of renal function and death ^47,48^. Therefore, we determined the kidney fungal burden 4 days post infection. Strikingly, no differences in renal fungal burden could be observed after 4 days of infection (**Fig. 1E**). We have previously shown that EphA2 activation triggers receptor-mediated invasion of oral epithelial cells ^33,40^. The mouse model of disseminated candidiasis leads to rapid organ dissemination and clearance of >99% of the fungus from the bloodstream within the first hour after intravenous injection ^49^, while *C. albicans* mutants defective in invasion have reduced kidney fungal burden ^50,51^. To rule out a possible decrease of dissemination out of the bloodstream we collected kidneys from WT and *Epha2*^−/−^ mice 12 hours post infection and enumerated fungal burden. Resistance and tolerance are two complementary host defense mechanisms that increase host fitness in response to invading *C. albicans* ^52^. Since WT and *Epha2*^−/−^ mice had similar renal fungal burden (**Fig. 1F**), we concluded that EphA2 deficiency enhances host tolerance, independent of fungal trafficking out of the bloodstream. Although EphA2 enhances neutrophilic killing of opsonized yeast ^34^, EphA2 has no effect on elimination of hyphae (**Fig. S2**), which are the dominant morphotype in kidneys after 12 hours of systemic infection (**Fig. S2**).

Severe renal failure plays a major role in lethality of systemic *C. albicans* infection ^9,53-55^. Therefore, we determined apoptotic areas in kidneys of WT and *Epha2*^−/−^ mice infected *C. albicans* using terminal deoxynucleotidyl transferase dUTP nick end labeling (TUNEL) staining ^48^. Although apoptosis could be detected in kidneys of both mouse strains, the overall apoptotic areas decreased in infected *Epha2*^−/−^ mice (**Fig. 1G; Fig S3**). Next, we assessed serum neutrophil gelatinase-associated lipocalin (NGAL), a marker of acute kidney injury ^48,56^. NGAL was strongly present in serum of WT mice, while this kidney injury marker was reduced in *Epha2*^−/−^ mice (**Fig. 1H**). During disseminated candidiasis, both systemic inflammation as well as rapid deterioration of the infected host resembles hyper-inflammatory sepsis ^57^. Therefore, we measured the sepsis marker soluble triggering receptor expressed on myeloid cells (TREM1) ^58^ in serum of infected mice. TREM1 was significantly reduced in *Epha2*^−/−^ mice compared to WT mice (**Fig. 1I**). Given that EphA2 promotes inflammation during oral *C. albicans* infection ^33,34,39-41^, we assessed the contribution of EphA2 in a mouse model of zymosan (β-glucan) - induced acute kidney injury (AKI), in which immune cells and inflammation exert essential roles in kidney damage ^59,60^. We found that EphA2 deficient mice were more resistant to AKI (**Fig. 1J**). Together, this data suggest that EphA2 promotes inflammation to accelerate disease progression during disseminated candidiasis.

### EphA2 promotes renal inflammation during disseminated candidiasis

Although the host immune response is required to control C. albicans infections, the inflammatory response causes significant collateral tissue damage. Therefore, we determined the cytokine and chemokine response in infected kidneys in WT and *Epha2*^−/−^ mice. *Epha2*^−/−^ mice had reduced kidney levels of several pro-inflammatory cytokines, including TNFα, IL-1β, and IL-6 (**Fig. 2A)**. By contrast, we found that *Epha2*^−/−^ mice had increased levels of IL-23, IFNγ, and IL-4 (**Fig. 2B**). Next, we assessed renal leukocyte infiltration during infection. *Epha2*^−/−^ mice had decreased accumulation of neutrophils and monocytes (**Fig. 2C; Fig. S4**), but no differences in macrophage accumulation was observed (**Fig. 2C**). Consistent with previous observations, EphA2 deficiency increased the accumulation of dendritic cells (DCs) during infection ^61^ (**Fig. 2D-F**). These experiments suggested that EphA2 is critical to promote renal inflammation during disseminated candidiasis.

**Figure 2.**
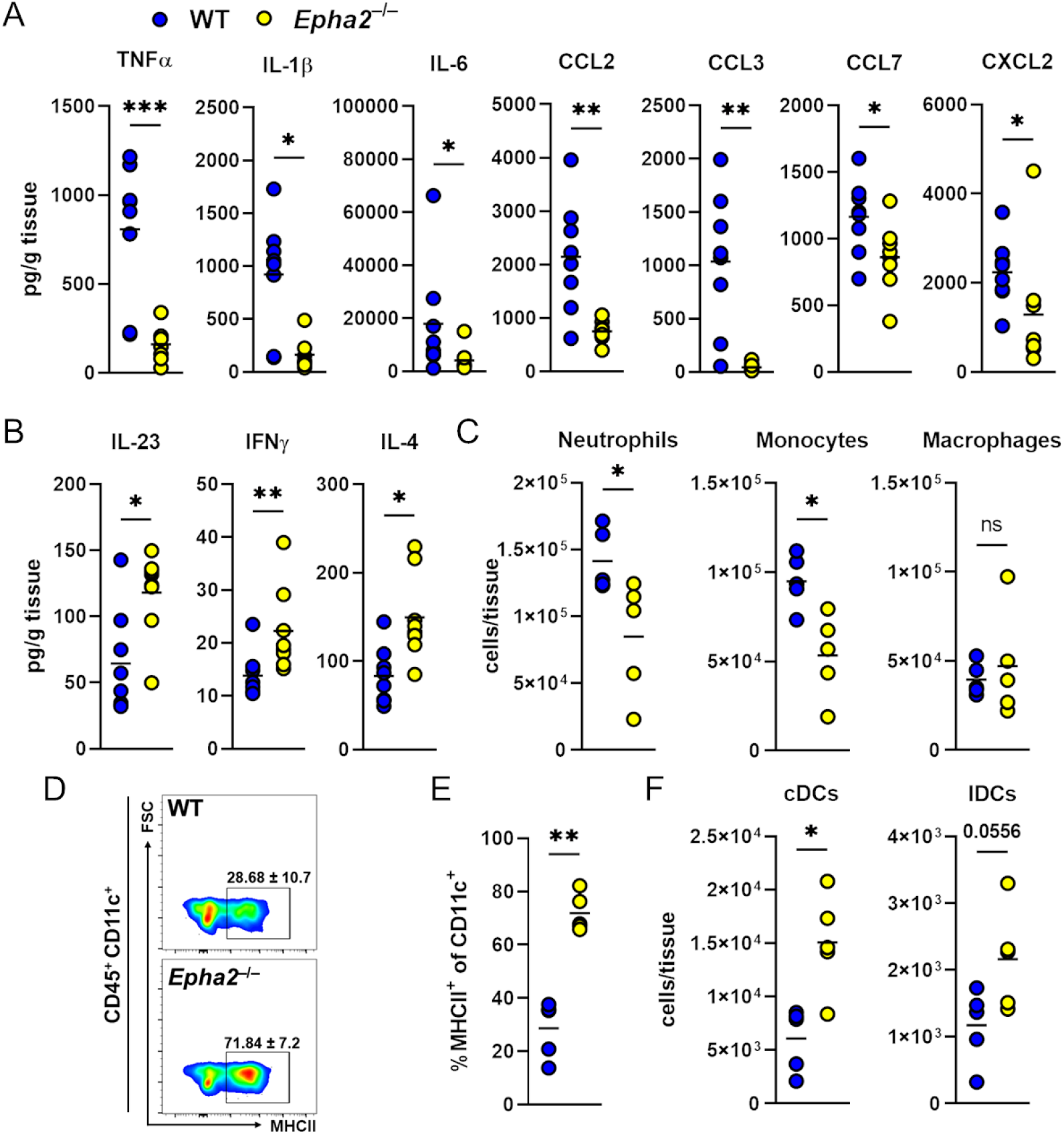
EphA2 deficiency reduces renal neutrophil and monocyte recruitment, but increases DC accumulation during candidiasis. **A**-**B** Level of indicated cytokines in kidneys after 3 days of infection. N=8; two independent experiments. Statistical significance is indicated by ^*^*P<*0.05; ^**^*P<*0.01; ^***^*P<*0.001. Mann-Whitney Test. **C** Accumulation of neutrophils, monocytes, and macrophages in the kidney of wild type and *Epha2*^−/−^ mice after 3 days of infection. N=5, two independent experiments. Statistical significance is indicated by; ^*^*P<*0.05. ns; No Significance. Mann-Whitney Test. **D**-**E** Representative flow cytometry plots of MHCII expression and frequencies of MHCII–expressing cells in infected kidneys after 3 days of infection. N=5; two experiments. **F** Accumulation of conventional dendritic cells (cDCs) and lymphoid dendritic cells (lDCs) in kidneys of wild type and *Epha2*^−/−^ mice after 3 days of infection (N=5). Statistical significance is indicated by ^*^*P<*0,05; ^**^*P<*0,01. Mann-Whitney Test.

### EphA2 deficiency reduces renal ferroptosis during disseminated candidiasis

To comprehensively evaluate the transcriptional response of *Epha2*^−/−^ mice during *C. albicans* infection, we performed RNA sequencing of infected kidneys. Consistent with our findings that EphA2 deficient mice had reduced renal apoptosis (**Fig. 1G**) and reduced inflammation (**Fig. 2**), KEGG pathway analysis revealed downregulation of genes involved in apoptosis and several immune response pathways (**Fig. 3A**). Using Gene Set Enrichment Analysis (GSEA) we found that genes involved in ferroptosis were significantly enriched in infected kidneys from WT mice compared to *Epha2*^−/−^ mice (**Fig. 3B**). In tumor cells, SLC7A11-mediated cystine uptake promotes GPX4 protein synthesis to reduce sensitivity to ferroptotic cell death ^62,63^. Therefore, we determined the two anti-ferroptotic genes *SLC7a11* and *GPX4*, as well as *LYZ2*, a myeloid cell lineage marker, using RNAscope (**Fig. 3C**). *SLC7a11* and *GPX4* RNAscope particles were reduced in infected kidneys from *Epha2*^−/−^ mice (**Fig. 3D**). Furthermore, *GPX4* particles in myeloid cells were enriched in kidney sections from WT compared *Epha2*^−/−^ mice (**Fig. 3E**). Using immunofluorescence, we confirmed that GPX4 protein expression was reduced in *Epha2*^−/−^ mice in infected tissue (**Fig. 3F**). Ferroptosis is a lipid peroxidation-driven form of RCD ^64^. Therefore, we determined the level of lipid peroxidation using 4-hydroxynonenal (4HNE) staining. While kidney sections of WT mice showed strong lipid peroxidation, sections of *Epha2*^−/−^ mice had low levels of ferroptosis (**Fig. 3G**) suggesting that during infection host cells undergo excessive lipid peroxidation although anti-ferroptotic mechanisms are upregulated. Since *LYZ2*^*–*^ cells in kidney sections from WT and *Epha2*^−/−^ mice had comparable *GPX4* expression, we tested whether EphA2 deficiency in macrophages (*LYZ2*^+^) results in decreased ferroptosis. Using the lipid peroxidation sensor BODIPY, as well as 4HNE staining, we showed that BMDMs from *Epha2*^−/−^ mice underwent similar magnitudes of ferroptosis compared to WT BMDMs (**Fig. S5**). Collectively, this data suggest that EphA2 promotes ferroptotic cell death via an extrinsic pathway during candidiasis.

**Figure 3.**
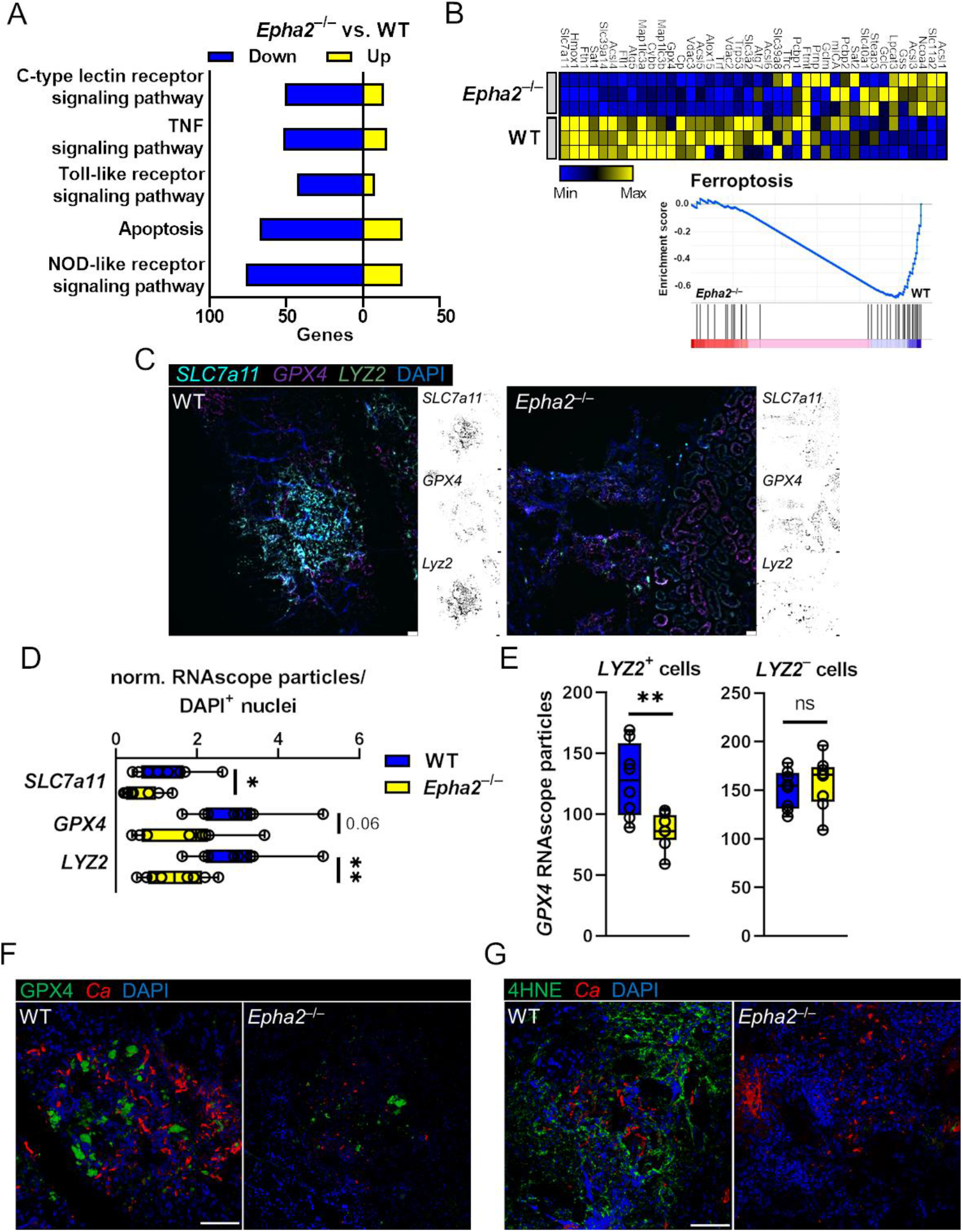
EphA2 promotes ferroptotic host cell death during candidiasis. **A** Up- and down regulated number of genes of KEGG pathways. RNASeq was performed on mRNA isolated from kidneys of WT and *Epha2*^™/™^ mice after 3 days of infection. N=3 per mouse strain. **B** GSEA of ferroptosis pathway genes. Heatmap of ferroptosis gene expression analysis in WT and *Epha2*^*-/-*^ mice. Normalized enrichment score is shown on Y-axis. **C** Representative RNAscope image of *SCL7a11, GPX4*, and *LYZ2* expression in infected kidneys after 3 days of infection. Scale bar 50 μm. **D-E** Quantification of RNAsope particles. (D) Normalized particles of DAPI^+^ nuclei. (E) *GPX4*^+^ particles of *LYZ2* positive and negative cells. N=4 per animal. **F** GPX4 protein expression in infected kidneys after 3 days of infection. GPX4 shown in green, *C. albicans* (*Ca*) in red. Tissue is visualized using DAPI. Scale bar 100 μm. **G** Lipid peroxidation in infected kidneys after 3 days of infection using 4HNE. 4HNE shown in green, *C. albicans* (*Ca*) in red. Tissue is visualized using DAPI. Scale bar 100 μm.

### Ferroptotic cell death exerts inflammation and promotes disease severity during candidiasis

To investigate whether ferroptotic cell death exacerbates inflammation and disease progression during fungal infection, we first determined that *C. albicans* induces ferroptosis in bone marrow-derived macrophages (BMDMs) (**Fig. 4A**) and renal tubular epithelial cells (RTECs) (**Fig. 4B**). Inhibition of ferroptosis using the selective inhibitor Ferrostatin-1 (Fer-1) ^65^ increased survival of BMDMs and RTECs during *C. albicans* interactions (**Fig. 4C, D; Fig. S6**). This finding is consistent with previous reports showing that Fer-1 treatment reduces macrophage cell death during *Histoplasma capsulatum* infection ^66^. Increased BMDM survival was associated with increased *C. albicans* killing (**Fig. 4E**). Furthermore, inhibition of ferroptosis reduced cytokine secretion in BMDMs (**Fig. 4F**) and RTECs (**Fig. 4G**) suggesting that ferroptosis exerts inflammation during *C. albicans* infection. To analyze the contribution of ferroptotic cell death to the pathogenicity of disseminated candidiasis, we infected WT mice with *C. albicans* followed by daily treatment with Fer-1. Mice treated with Fer-1 were less susceptible to *C. albicans* challenge compared to vehicle control mice (**Fig. 4H, I**). Collectively, disseminated candidiasis induces ferroptosis in various host cell types to promote inflammation and disease progression.

**Figure 4.**
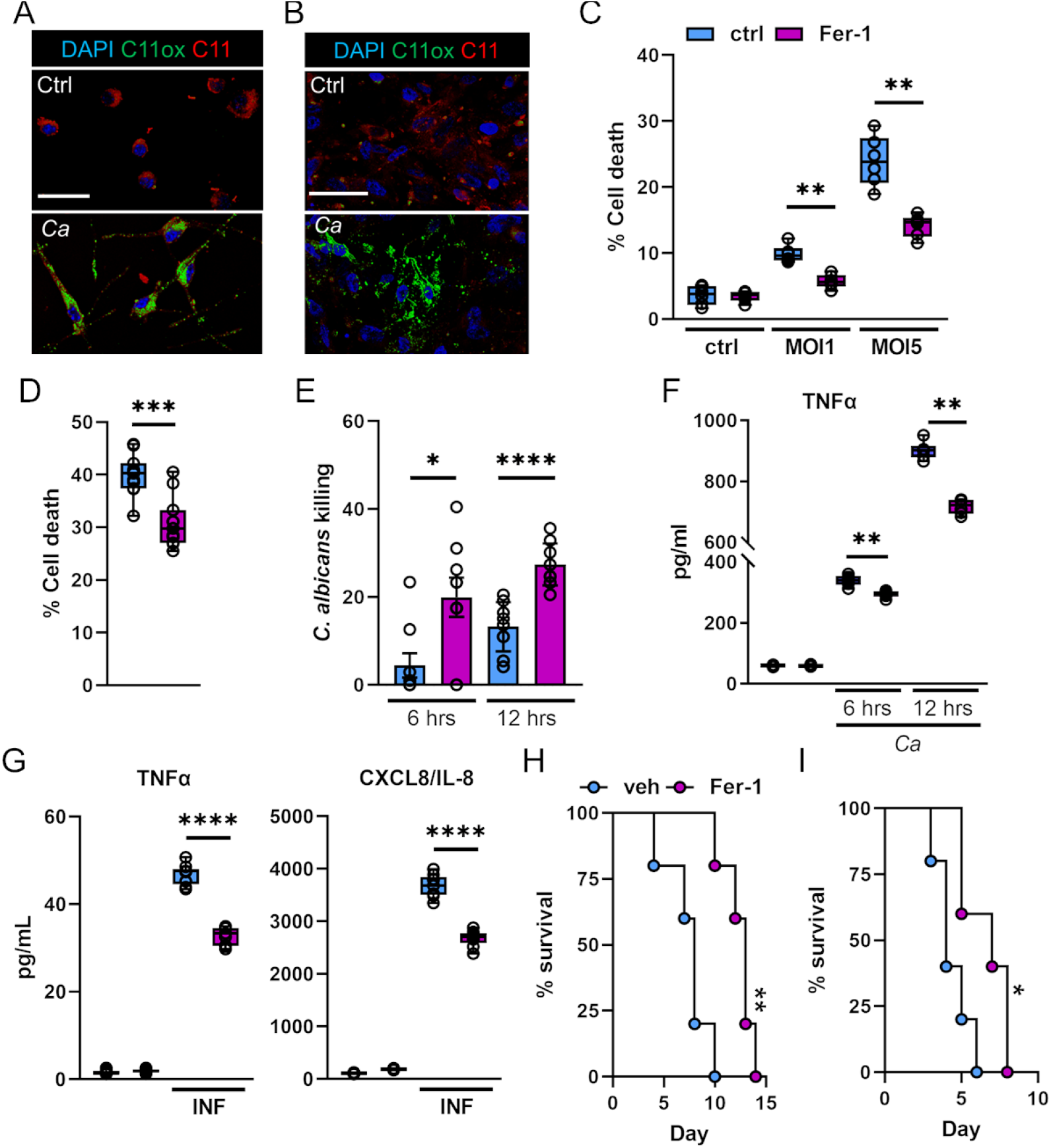
Ferroptotic cell death exerts inflammation, decreases macrophage-mediated fungal killing, and promotes disease progression during candidiasis. **A-B** Representative images of C11 oxidation of BM-derived macrophages (A) and renal tubular epithelial damage (B) during *C. albicans* infection. Host cells were infected with an MOI of 1 (BMDMs, 4 hours; RTECs, 2 hours). **C** BM-derived macrophages were treated with 10μM Fer-1 for 1 hour followed by interaction with *C. albicans* (MOI 1 and 5) for 4 hours. PI^+^ (dead) cells were determined by gating on F4/80^+^ cells. N=3; duplicate. ^**^*P<*0.01; Mann-Whitney Test. **D** Renal tubular epithelial damage determined by specific ^51^Cr release. N=3; triplicate. ^***^*P<*0.001; Mann-Whitney Test **E** *C. albicans* killing of macrophages treated with Fer-1. MOI 1. N=3; triplicate. ^*^*P<*0.05; ^***^*P<*0.001; Mann-Whitney Test. **F** TNFα secretion of BM-derived macrophages during *C. albicans* infection (MOI 1) for indicated time points. N=3, duplicate. ^**^*P<*0.01; Mann-Whitney Test. **G** CXCL8 and TNFα secretion of RTECs in the presence and absence of Fer-1 during C. albicans infection. MOI 5. ^****^*P<*0.0001; Mann-Whitney Test. **H-I** Survival of WT mice treated daily with 10 mg/kg Fer-1 or vehicle control. Inoculum 1.25×10^5^ (H) and 2.5×10^5^ (I) *C. albicans*. N=5; two independent experiments per inoculum. ^*^*P <* 0.05; ^**^*P <* 0.01. Mantel-Cox Log-Rank test.

### EphA2 and JAK signaling limit IL-23 secretion in DCs

Following systemic *C. albicans* infection, DCs produce IL-23 to stimulate natural killer (NK) cell activity ^8^. Accordingly, renal IL-23 levels correlated with increased cDC infiltration during infection in *Epha2*^−/−^ mice (**Fig. 5A**). DCs derived from *Epha2*^−/−^ BM secreted more IL-23 when stimulated with β-glucan (**Fig. 5B**), while TNFα levels were unaffected (**Fig. S7**). Next, we examined the transcriptional response of β-glucan stimulated DCs using RNA sequencing. KEGG pathway mapping revealed downregulation of genes associated with Janus-associated kinase (JAK)-Signal transducers and activators of transcription (STAT) signaling in *Epha2*^−/−^ BMDCs, while genes of the peroxisome proliferator-activated receptors (PPAR) pathway were upregulated (**Fig. 5C**). To investigate the contribution of JAK-STAT and PPAR signaling to IL-23 secretion in DCs, we treated WT BMDCs with Ruxolitinib (JAK1/2 inhibitor), the PPARγ antagonist GW9662, and the PPARγ agonist Rosiglitazone followed by β-glucan stimulation. While PPARγ stimulation or inhibition had no effect on IL-23 secretion, Ruxolitinib increased IL-23 levels in supernatants during β-glucan stimulation (**Fig. 5D**). This experiment suggested that β-glucan-induced JAK-STAT signaling reduces IL-23 secretion in DCs.

**Figure 5.**
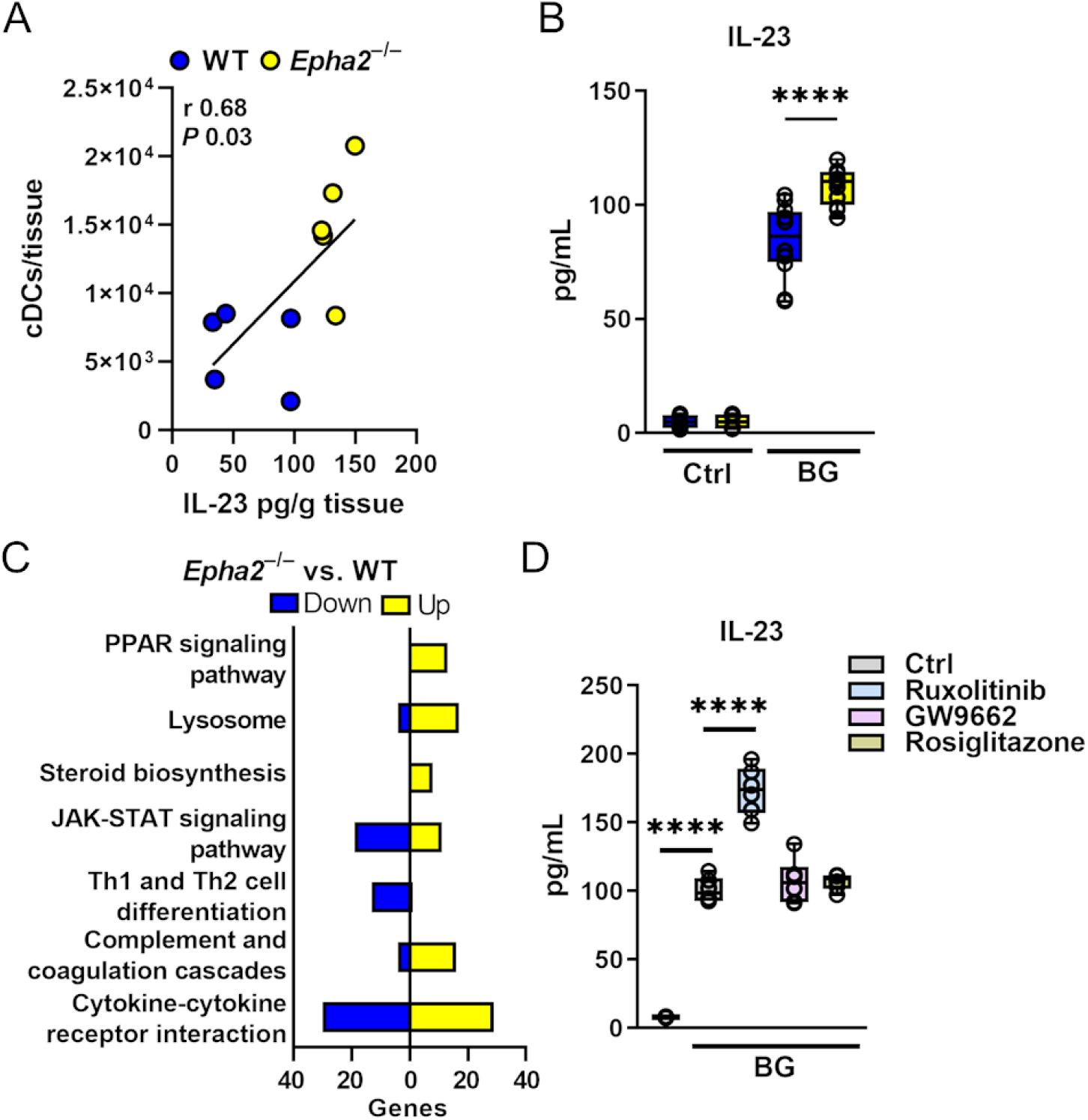
EphA2 signaling decreases IL-23 secretion in BMDCs. **A** Correlation of IL-23 kidney levels and cDC infiltration during candidiasis in WT and *Epha2*^−/−^ mice. N=5. Pearson correlation coefficient ^*^, *P <* 0.05. **B** IL-23 levels in supernatants of DCs after 24 hours stimulation with β-glucan (curdlan). DCs generated from WT and *Epha2*^−/−^ mice. N=6, duplicate. ^****^*P<*0.0001, Mann-Whitney Test. Ctrl, control; BG, β-glucan; *Ca, C. albicans*. **C** Number of up and down regulated genes in corresponding KEGG pathways. BMDCs from WT and Epha2–/– mice (n=3) were stimulated for 6 hours with 25 μg/mL curdlan. **D** DC IL-23 secretion treated with indicated inhibitors and stimulated with curdlan. 24 hours post stimulation IL-23 levels in supernatants were measured with ELISA. N=3; duplicate. ^****^*P<*0.0001, Mann-Whitney Test.

### IL-23 inhibits ferroptosis during disseminated candidiasis

Besides stimulating NK cells ^8^, IL-23 secures myeloid cell survival during candidiasis ^67^. It is thought that this mechanism is key for maintaining sufficient numbers of phagocytes at the site of infection to ensure efficient host protection ^67^. The link between myeloid cell survival and IL-23 was intriguing since IL-23 receptor downstream targets have been associated to counteract ferroptotic cell death ^68,69^. To test if IL-23 prevents macrophage ferroptosis during *C. albicans* infection, we measured total cell fluorescence of oxidized C11-BODIPY. Exogenous IL-23 reduced macrophage lipid peroxidation during *C. albicans* interaction (**Fig. 6A**). Furthermore, treatment with IL-23 increased macrophage survival (**Fig. 6B**), their *C. albicans* killing capacity (**Fig. 6C**), and reduced inflammation (**Fig. 6C**), as seen in macrophages treated with the ferroptosis inhibitor (**Fig. 4**). This data suggested that IL-23 signaling reduces ferroptotic macrophage cell death during *C. albicans* infection. Next, we treated *Candida*-infected WT mice with recombinant murine IL-23 for 3 consecutive days (**Fig. 6D**). IL-23 treatment increased the median survival by >65% (6 vs. 10 days) (**Fig. 6D**) and reduced the renal fungal burden (**Fig. 6E**). We determined ferroptosis in infected kidneys by staining section for 4HNE. Treatment with rmIL-23 decreased lipid peroxidation compared to PBS control (**Fig. 6G**). Collectively, we show that IL-23 signaling prevents inflammatory ferroptosis in myeloid cells and improves disease outcomes during disseminated candidiasis.

**Figure 6.**
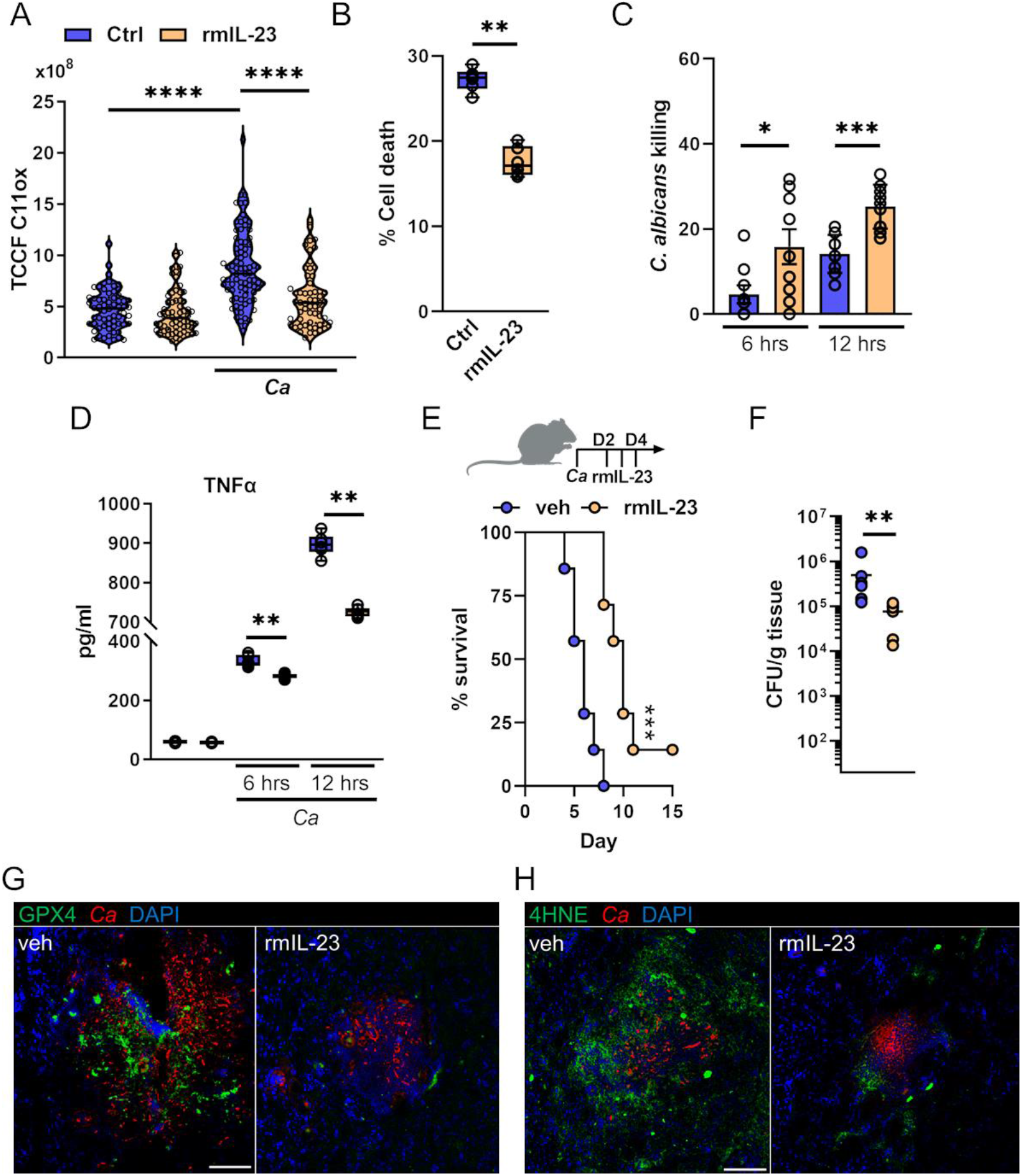
IL-23 signaling reduces ferroptosis during *C. albicans* infection. **A** BM-derived macrophages were infected with C. albicans (MOI 1) for 4 hours in the presence and absence of rmIL-23. Total C11ox fluorescence was quantified. Individual cells (N=57-80; 3 independent experiments) were identified and total fluorescence was measured using ImageJ. ^****^*P<*0.0001, Mann-Whitney Test. **B** BM-derived macrophages were treated with rmIL-23 for 1 hour followed by interaction with *C. albicans* (MOI 5) for 4 hours. PI^+^ (dead) cells were determined by gating on F4/80^+^ cells. N=3; duplicate. ^**^*P<*0.01; Mann-Whitney Test. **C** Macrophages-mediated *C. albicans* killing in the presence and absence of rmIL-23. MOI 1. N=3; triplicate. ^*^*P<*0.05; ^***^*P<*0.001; Mann-Whitney Test **D** TNFα secretion of BM-derived macrophages during *C. albicans* infection (MOI 1) for indicated time points. N=3, duplicate. ^**^*P<*0.01; Mann-Whitney Test. **E** (Top) IL-23 treatment scheme during disseminated candidiasis. (Bottom) Survival of wild-type mice infected intravenously with 2.5×10^5^ SC5314 *C. albicans*. N=7; two independent experiments. Mice were treated with recombinant murine IL-23 (rmIL-23) or PBS. Statistical significance is indicated by ^***^, *P <* 0.001. Mantel-Cox Log-Rank test. **F** Renal fungal burden after 3 days of infection. Single rmIL-23 treatment at day 2. N=6-7; two independent experiments. Mice were treated with recombinant murine IL-23 (rmIL-23) or vehicle (veh; PBS). ^**^*P<*0.01, Mann-Whitney Test. **G** GPX4 protein expression in infected kidneys after 3 days of infection. Single rmIL-23 treatment at day 2 relative to infection. GPX4 shown in green, *C. albicans* (*Ca*) in red. Tissue is visualized using DAPI. Scale bar 100 μm. **H** Lipid peroxidation in infected kidneys after 3 days of infection using 4HNE. Single rmIL-23 treatment at day 2 relative to infection. 4HNE shown in green, *C. albicans* (*Ca*) in red. Tissue is visualized using DAPI. Scale bar 100 μm.

## Discussion

Distinct RCD mechanisms promote the resolution of infection by destroying intracellular niches which benefit the pathogen, and to coordinate an appropriate innate immune response thereafter ^23^. However, the RCD ferroptosis has been implicated in the development of many diseases ^70-72^. Although inflammation is required to fight infections, a reduction in myeloid cell-mediated immunopathology may lead to pathogen tolerance, a phenomenon whereby the host is able to better resist infection by reducing tissue damage ^18,73^. Our findings uncover a mechanism linking ferroptosis to immunopathology during candidiasis (**Fig. S8**). Here we show that myeloid and stromal cells undergo ferroptotic RCD to accelerate inflammation during fungal encounter. Macrophage ferroptosis is limited by exogenous IL-23; thus safeguarding efficient fungal clearance and controlled inflammation.

While the production of reactive oxygen species (ROS) is a key aspect of phagocyte-mediated host responses during *C. albicans* infection, an increase in ROS may cause lipid peroxidation and ferroptosis ^74^. Here, we show that ferroptotic cell death benefits fungi by reducing the killing capacity of macrophages. Lipid peroxidation during ferroptosis results in production of 4HNE and malondialdehyde (MDA), which are able to react with primary amines on proteins or DNA to form crosslinks ^75^. Exogenous 4HNE impairs the PKC signaling pathway in RAW macrophages ^76^, a critical mediator of antifungal host defense ^77^. The contribution of lipid peroxides and their degradation products to immune signaling and antifungal defense both, intrinsic and extrinsic, is being further investigated.

In response to reduced inflammatory signals, classical macrophages (M1 state) are more resistant to pro-ferroptotic treatment with a specific GPX4 inhibitor ^78^. However, sensitization to ferroptotic cell death is regulated by various mechanical stimuli ^79^. Protrusive force is generated during phagocytosis ^80^, and immune cells respond to mechanical forces during the polarized growth of fungal hyphae ^81^. Likewise, fungal β-glucan recognition induces mTOR signaling in monocytes ^82^, which has been associated with increased sensitivity to ferroptosis ^83^. Hence, mechanical forces during infection and consequently activation of distinct downstream signaling pathways sensitize myeloid cells to undergo ferroptotic cell death.

IL-23 has received significant interest as a therapeutic target for a number of autoimmune conditions in recent clinical trials ^84^. However, IL-23 has established roles during antifungal immunity ^85^. IL-23 expression is strongly induced in response to *C. albicans* via the C-type lectin and TLR pathways ^86^ and is best known for regulating IL-17 production by T cells and innate lymphoid cells at epithelial barriers ^84^. IL-23 binds to IL-12Rβ1 and IL-23R followed by receptor complex signaling via JAK2 and Tyk2 ^85,87,88^. In a mouse model of disseminated candidiasis, pre-treatment with tofacitinib ^89^ and ruxolitinib ^90^ (JAK inhibitors) increase susceptibility and fungal burden, while inflammation increases when therapy is started at the onset of disease ^90^. Furthermore, the IL-23 signaling cascade activates several STAT members, including STAT3 ^91,92^. In tumor cells, STAT3 inhibition induces ferroptosis via Nrf2-GPX4 signaling ^68^, while STAT3 activation suppresses expression of ACSL4, an enzyme that enriches membranes with long polyunsaturated fatty acids and is required for ferroptosis ^69^. Nur *et al*. showed that IL-23 secures survival of myeloid cells during candidiasis by inhibiting apoptosis ^67^. Here, we demonstrate a non-canonical role of IL-23 signaling in inhibiting macrophage ferroptosis during fungal infection. Historically, cell death pathways have long been considered to function in parallel with little or no overlap. However, it is currently known that lytic and non-lytic RCDs, such as apoptosis, necroptosis, pyroptosis, and ferroptosis are tightly connected, and can cross-regulate each other ^22^. For instance STAT3 activation (downstream of IL-23R) limits ferroptosis, pyroptosis, and necroptosis ^68,93,94^. This suggest that different types of cell death depend on the stimulant, either infectious agent or drug, the cell type, and the environment, but share similar downstream signals and molecular regulators.

While IL-23 inhibits ferroptotic cell death in macrophages, some cytokines, such as INFγ, drive ferroptosis via STAT1 signaling in tumor cells ^95^. Whether the inflammatory tissue environment during infection and their corresponding cytokines sensitize immune cells to ferroptotic cell death needs to be determined. Although ferroptotic cells exhibit phosphatidylserine surface exposure, which triggers removal of dying cells by macrophages ^96^, ferroptotic cells are poorly engulfed and cleared ^97^. Thus, we speculate that inefficient clearance of ferroptotic immune cells might accelerate immunopathology during infection.

In DCs β-glucan recognition induces several signaling pathways, including AKT, MAPKs, IKK, and NF-κB ^86^. Here we show that activation of the β-glucan receptor EphA2 represses IL-23 secretion suggesting that β-glucan recognition stimulates IL-23 expression via Dectin-1/TLR-2 ^98^, but limits the cytokine secretion via non-classical β-glucan recognition. EphA2 activates JAK1/STAT3 signaling ^99^, and STAT3 deficient DCs exhibit increased IL-23 production after stimulation ^100^. Accordingly, EphA2 deficient DCs and inhibition of JAK signaling in β-glucan stimulated DCs increase IL-23 secretion. Taken together, EphA2-JAK-STAT signaling negatively regulates an inflammatory IL-23 DC phenotype during fungal encounter Collectively, our study demonstrates that ferroptotic host cell death is linked to immunopathology and can be targeted by recombinant cytokine therapy during fungal infection. We postulate that strategies to inhibit ferroptotic cell death during infection will have important therapeutic benefits.

## Methods

### Ethics statement

All animal work was approved by the Institutional Animal Care and Use Committee (IACUC) of the Lundquist Institute at Harbor-UCLA Medical Center.

### Subject details

For *in vivo* animal studies, age-and sex matched mice were used. Animals were bred/housed under pathogen-free conditions at the Lundquist Institute. Animals were randomly assigned to the different treatment groups. Researchers were not blinded to the experimental groups because the endpoints (survival, fungal burden, cytokine levels) were objective measures of disease severity. *Epha2*^−/−^ (B6-Epha2^*tm1Jrui*^/J) mice were provided by A. Wayne Orr ^46^. C57BL/6 control and *Clec7a*^−/−^ mice were purchased from The Jackson Laboratory. All mice were cohoused for at least 1 week before the experiments.

### Mouse model of HDC

Resistance to disseminated candidiasis was tested in the mouse model of HDC using 6- to 8-week-old mice (C57BL/6J background) as previously described ^17^. The *C. albicans* SC5314 strain was serially passaged 3 times in YPD broth, grown at 30°C at 200 rpm for 16–24 hours at each passage. Yeast cells were washed, and 2.5× 10^5^ or 1.25×10^5^ *C. albicans* cells injected intravenously via the lateral tail vein. For survival experiments, mice were monitored three times daily and moribund mice were humanely euthanized. To determine organ fungal burden, mice were sacrificed after 12 hours and 4 days of infection, after which the kidneys were harvested, weighed, homogenized, and quantified on sabouraud dextrose agar plates containing 80 mg/L chloramphenicol. For histology, mouse kidneys were fixed in 10% buffered formalin and embedded in paraffin.

To inhibited ferroptosis during infection mice were treated daily intraperitoneally (start 6 hours post infection) with 10 mg/kg Ferrostatin-1 (SelleckChem; >99% purity) dissolved in 0.9% NaCl. In another experiment mice were treated intravenously 2 days post infection with 12.5 μg/kg recombinant murine IL-23 (1887-ML-010/CF, R&D Systems) for 3 consecutive days.

### Immunohistochemistry

Apoptotic cell death was determined as previously described ^48^. Briefly, terminal deoxynucleotidyl transferase nick end labeling (TUNEL) staining was performed using the *in situ* apoptosis detection kit (ApopTag, S7100, Chemicon, Temecula, CA, USA) according to the manufacturer’s protocol with minor modifications. The paraffin-embedded renal sections were placed on poly-L-lysine coated glass slides, deparaffinized in xylene and rehydrated in a graded series of alcohol. Then treated with protease K (20 g/ml) for 15 min at room temperature. Sections were incubated with reaction buffer containing terminal deoxynucleotidyltransferase at 37°C for 1 h. After washing with stop/wash buffer, sections were treated with anti-digoxigenin conjugate for 30 min at room temperature and subsequently developed color in peroxidase substrate. The nuclei were counterstained with 0.5% methyl green. TUNEL-positive cells/areas were determined by bright field microscopy. For quantification, apoptotic areas were quantified using PROGRES GRYPHAX® software (Jenoptik).

To determine GPX4 and 4HNE expression, kidneys of WT and *Epha2*^−/−^ mice were harvested 3 days post infection, and snap frozen in Tissue-Tek® OCT. 10 μm kidney sections were fixed with cold acetone, rehydrated in PBS, blocked with BSA, and stained overnight using anti-GPX4 or anti-4HNE (ab125066 and ab46545, respectively). Sections were washed and incubated with anti-rabbit IgG (H+L) coupled with Alexa Fluor™ 488 (Thermo Fisher Scientific). *C. albicans* was detected with an anti-*Candida* antiserum (Biodesign International) conjugated with Alexa Fluor 568 (Thermo Fisher Scientific). To visualize nuclei, cells were stained with DAPI (Prolong Gold antifade reagent with DAPI). Images of the sections (z-stack) were acquired with a Leica TCS SP8 confocal microscope. To enable comparison of the fluorescence intensities among slide, the same image acquisition settings were used for each experiment.

### Determination of serum NGAL and TREM-1

Serum NGAL and TREM-1 were measured at day 3 post-infection. Blood was collected by cardiac puncture from each mouse at the time of sacrifice, and stored at -80°C until use. NGAL and TREM-1 concentrations were determined using DuoSet ELISA Kit (DY1857-05 & DY1187, R&D Systems).

### Cytokine and chemokine measurements *in vivo*

To determine the whole kidney cytokine and chemokine protein concentrations, the mice were intravenously infected with *C. albicans* SC5314. The mice were sacrificed after 3 days post-infection, and their kidney were harvested, weighed and homogenized. The homogenates were cleared by centrifugation and the concentration of inflammatory mediators was measured using the Luminex multiplex bead assay (Invitrogen).

### Generation of BM chimeric mice

Bone marrow chimeric mice were generated as previously described ^34^. Briefly, for BM cell transfers, femurs and tibias from 6- to 8-week-old donor wild-type (*Epha2*^+/+^; CD45.1, or CD45.2) and *Epha2*^−/−^ (CD45.2) mice were removed aseptically and BM was flushed using cold PBS supplemented with 2 mM EDTA. Recipient wild-type (CD45.1; B6.SJL-*Ptprc*^*a*^ *Pepc*^*b*^/BoyJ) and *Epha2*^−/−^ mice were irradiated with 10 Gy and were reconstituted 6 hours after irradiation with 2.5×10^6^ *Epha2*^+/+^ CD45.2 (WT→WT), *Epha2*^+/+^ CD45.1 (wild-type *Epha2*^−/−^), or *Epha2*^−/−^ CD45.2 BM cells by lateral tail-vein injection. Mice were given enrofloxacin (Victor Medical) in the drinking water for the first 4 weeks of reconstitution before being switched to antibiotic-free water. Chimeras were infected with *C. albicans* 10 weeks after transplantation. Prior to infection, we confirmed that mice reconstituted with congenic BM stem cells had achieved a satisfactory level of chimerism by assessing the number of CD45.1 and CD45.2 leukocytes in the blood, using flow cytometry (**Fig. S1**).

### Flow cytometry of infiltrating leukocytes

Immune cells in the mouse kidney were characterized as described. Briefly, mice were infected with *C. albicans* strain SC5314. After 3 days of infection, mice were anesthetized using ketamine/xylazine and perfused with 10 ml of PBS before kidney harvesting. Single-cell suspensions from kidney were prepared using previously described methods ^101^. In brief, kidneys were finely minced and digested at 37°C in digestion solution (RPMI 1640 with 20 mM HEPES [Gibco] without serum) containing liberase TL (Roche) and grade II DNAse I (Roche) for 20 minutes with shaking. Digested tissue was passed through a 70-μm filter and washed. The remaining red blood cells were lysed with ACK lysis buffer (Lonza). The cells were suspended in 40% Percoll (GE Healthcare). The suspension was overlaid on 70% Percoll and centrifuged at 836 g for 30 minutes at room temperature. The leukocytes and nonhematopoietic cells at the interphase were isolated, washed 3 times in FACS buffer (0.5% BSA and 0.01% NaN3 in PBS). The single-cell suspensions were then incubated with rat anti-mouse CD16/32 (2.4G2; BD Biosciences) for 10 minutes (1:100) in FACS buffer at 4°C to block Fc receptors. For staining of surface antigens, cells were incubated with fluorochrome-conjugated (BV421, BV711, FITC, PE, PE-Cy7, allophycocyanin [APC], APC-Cy7) antibodies against mouse CD45 (30-F11; BD Biosciences), Ly6C (AL-21; BD Biosciences), Ly6G (1A8, BioLegend), CD11b (M1/70; eBioscience), CD11c (N41, BioLegend), MHCII (M5/114.15.2, BioLegend), and CD206 (C068C2; BioLegend). After washing with FACS buffer, the cell suspension was stained with a Fixable Viability Stain 510 (BD Biosciences), washed, and resuspended in FACS buffer. The stained cells were analyzed on a FACSymphony system (BD Biosciences), and the data were analyzed using FACS Diva (BD Biosciences) and FlowJo software (Treestar). Only single cells were analyzed, and cell numbers were quantified using PE-conjugated fluorescent counting beads (Spherotech).

### RNA Sequencing

Total RNA was isolated as described before ^102^. Briefly, kidneys from infected mice were harvested at 3 days post infection andplaced for 1 hour in RNAlater solution (Invitrogen). Kidneys were homogenized in Lysing Matrix C tubes (MPbio), following RNA extraction with RNeasy (Qiagen). For RNA sequencing of BMDCs, cells were purified using negative magnetic bead selection (MojoSort Mouse Pan Dendritic Cell Isolation Kit, BioLegend). 2.5×10^6^ BMDCs per well were cultured for 6h in 6-well plates in RPMI complete (R10; 10% HI-FBS, 2mM L-glutamine, 100U/ml Penicillin, 100µ/ml Streptomycin, 50µM β-ME) and 20mg/ml GM-CSF in presence of 25µg/ml of Curdlan. RNA was isolated using the RNeasy Kit (Qiagen). RNA sequencing was performed by Novogene Corporation Inc. (Sacramento, USA). mRNA was purified from total RNA using poly-T oligo-attached magnetic beads. To generate the cDNA library the first cDNA strand was synthesized using random hexamer primer and M-MuLV Reverse Transcriptase (RNase H^™^). Second strand cDNA synthesis was subsequently performed using DNA Polymerase I and RNase H. Double-stranded cDNA was purified using AMPure XP beads and remaining overhangs of the purified double-stranded cDNA were converted into blunt ends via exonuclease/polymerase. After 3’ end adenylation a NEBNext Adaptor with hairpin loop structure was ligated to prepare for hybridization. In order to select cDNA fragments of 150∼200 bp in length, the library fragments were purified with the AMPure XP system (Beckman Coulter, Beverly, USA). Finally, PCR amplification was performed and PCR products were purified using AMPure XP beads. The samples were read on an Illumina NovaSeq 6000 with ≥20 million read pair per sample.

### Downstream Data Processing

Downstream analysis was performed using a combination of programs including STAR, HTseq, and Cufflink. Alignments were parsed using Tophat and differential expressions were determined through DESeq2. KEGG enrichment was implemented by the ClusterProfiler. Gene fusion and difference of alternative splicing event were detected by Star-fusion and rMATS. The reference genome of *Mus musculus* (GRCm38/mm10) and gene model annotation files were downloaded from NCBI/UCSC/Ensembl. Indexes of the reference genome was built using STAR and paired-end clean reads were aligned to the reference genome using STAR (v2.5). HTSeq v0.6.1 was used to count the read numbers mapped of each gene. The FPKM of each gene was calculated based on the length of the gene and reads count mapped to it. FPKM, Reads Per Kilobase of exon model per Million mapped reads, considers the effect of sequencing depth and gene length for the reads count at the same time. Differential expression analysis was performed using the DESeq2 R package (2_1.6.3). The resulting *P*-values were adjusted using the Benjamini and Hochberg’s approach for controlling the False Discovery Rate (FDR). Genes with an adjusted *P*-value < 0.05 found by DESeq2 were assigned as differentially expressed. To allow for log adjustment, genes with 0 FPKM are assigned a value of 0.001. Correlation were determined using the cor.test function in R with options set alternative = “greater” and method = “Spearman.” To identify the correlation between the differences, we clustered different samples using expression level FPKM to see the correlation using hierarchical clustering distance method with the function of heatmap, SOM (Self-organization mapping) and kmeans using silhouette coefficient to adapt the optimal classification with default parameter in R. We used clusterProfiler R package to test the statistical enrichment of differential expression genes in KEGG pathways. The high-throughput sequencing data from this study have been deposited with links to BioProject accession number PRJNA773053 and PRJNA773073 in the NCBI BioProject database.

### RNAscope

*SCL7a11, Gpx4 and Lyz2* mRNA was detected using RNAscope® *in situ* hybridization. Sections were thawed and postfixed for 15 min in 4% paraformaldehyde (4C°), and washed in PBS at room temperature. *In situ* hybridization was performed using the RNAscope® Multiplex v2 Fluorescent Assay (Advanced Cell Diagnostics, Inc.) in strict accordance with the manufacturer’s instruction. Probes used were against mouse *Slc7a11* (RNAscope® Probe-Mm-C1, Mm-Slc7a11), *Gpx4* (RNAscope® Probe-Mm-C2, Mm-Gpx4-O1 targeting 12-877 of NM_008162.4) and Lyz2 (RNAscope® Probe-Mm-C3, Mm-Lyz2). The commercially available negative control probe was used, which is designed to target the *DapB* gene from *Bacillus subtilis*. In brief, endogenous peroxidases present in the tissue were blocked with an RNAscope® hydrogen peroxidase solution. Tissue was washed in distilled water, then immersed in 100% ethanol, air dried and a hydrophobic barrier was applied to the slides. The sections were permeabilized with RNAscope® protease III for 30 min at 40°C. Sections were hybridized with the *Slc7a11, Gpx4, and Lyz2* probes at 40°C for 2 h. This was followed by a series of amplification incubation steps: Amp 1, 30 min at 40°C; Amp 2, 30 min at 40°C; Amp 3, 15 min at 40°C. Sections were washed with provided washing buffer 2 × 2 min in between each amplification step. Assignment of Akoya 520 (FITC slc7a11), 570 (Cy3 Gpx4) and 690 (Cy5 Lyz2) occurred with a peroxidase blocking step sequentially. Finally, DAPI stain was applied and sections were coverslipped with Invitrogen Prolong Gold antifade mounting medium. Images were taken with the Leica 3D culture Thunder imaging system. For analysis of two representative sections of the renal cortex were taken for each section for two sections (total of n=4 per animal). Images were analyzed using Image J. Each channel was thresholded with the Otsu filter circularity set to 0.25-1 and particle size set to 10-Infinity. Thresholded images were then analyzed and total counts were used to represent each gene. Each gene was then normalized to total cell count as assessed by DAPI positive cell count.

### Bone marrow isolation and cell differentiation

Bone marrow cells were flushed from femurs and tibias using RPMI 1640 medium (Gibco) supplemented with 10% HI-FBS and passed through a 70 µm cell strainer. Bone marrow derived macrophages were generated by growing freshly isolated bone marrow cells from WT and *Epha2*^−/−^ mice in presence of 25ng/ml M-CSF during 7 days in DMEM supplemented with 10% HI-FBS and Pen/Strep (100U/ml and 100µ/ml respectively, Gemini Bioproducts) at 37°C in a humidified atmosphere containing 5% CO_2_. Bone marrow derived dendritic cells were generated by growing freshly isolated bone marrow cells from WT and *Epha2*^−/−^ mice in presence of 20ng/ml GM-CSF during 8 days in RPMI complete (R10) at 37°C in a humidified atmosphere containing 5% CO_2_. BMDC purity was determined by flow cytometry (>95%).

### Neutrophil killing assay

The capacity of neutrophils to kill *C. albicans* hyphae was determined using the alamarBlue (Invitrogen) reduction as a measure of fungal inactivation. Neutrophils from mice were isolated as described above. Neutrophils were incubated in duplicate wells of flat bottom 96-well plates containing hyphae that had been grown for 3 hours with or without serum opsonization (2% heat-inactivated mouse serum; Gemini Bio-Products), at a neutrophil to *C. albicans* ratio of 1:4 at 37°C. After 2.5 hours, the neutrophils were lysed with 0.02% Triton X-100 in water for 5 minutes, after which the *C. albicans* hyphae were washed twice with PBS and incubated with 1 × alamarBlue (Invitrogen) for 18 hours at 37°C. Optical density at a wavelength of 570nm and 600nm was determined. Neutrophil killing capacity was calculated as the amount of alamarBlue reduced by wells containing *C. albicans* hyphae incubated with and without neutrophils.

### Macrophage killing assay

Bone marrow-derived macrophages were generated as described above. Macrophage killing of *C. albicans* yeast was determined by CFU enumeration. Briefly, unopsonized *C. albicans* SC5314 yeast cells were incubated with BMDMs in a 1:1 ratio for 6 and 12 hours, respectively. Macrophages were lysed with 0.02% Triton X-100 in ice-cold water for 5 minutes, diluted and remaining *Candida* cells were quantitatively cultured. To determine the effect of ferroptosis inhibition during killing BMDMs were incubated with 10 μM Fer-1 or 50ng/ml rmIL-23 for 1 hour prior to infection.

### Quantification of ferroptosis *in vitro*

After 7 days of differentiation BMDMs were collected and seeded on fibronectin-coated glass coverslips in presence or absence of 10µM Fer-1 or 50ng/ml rmIL-23, after 1h BMDMs were infected with *C. albicans* SC5314 at MOI 1. After 210 min Bodipy™ 581/591 C11 was added at the concentration of 10µM for 30 minutes. Cells were fixed using 2% paraformaldehyde diluted in PBS. After washing the coverslip with PBS, cells were mounted on microscopic glass using ProLong Gold Antifade Mountant with DAPI. The total fluorescent integrated density of the reduced form of Bodipy 581/591 C11 was quantified using ImageJ (v1.8). To measure individual cellular areas and mean fluorescence, an outline was drawn around each cell, along with several adjacent background readings. Total corrected cellular fluorescence (TCCF) was calculated using the following formula. Integrated density - (area of selected cell × mean fluorescence of background readings).

In another experiment we quantified 4HNE during macrophage-*C. albicans* interactions. After 240 min cells were fixed, washed, and permebalized following staining with anti-4HNE (ab46545; Abcam). TCFF was of 4HNE was determined as described above.

### Macrophage survival *in vitro*

1×10^6^ BMDMs seeded in 24 well plates were incubated with C. albicans (MOI 1 and 5, respectively) in presence or absence of 10µM Fer-1 or 50ng/ml IL-23. After 4 hours BMDMs were harvested and stained with an antibody against F4/80 (BM8, Biolegend) and propidium iodide (BD Biosciences). The stained cells were analyzed on a FACSymphony system (BD Biosciences).

### Renal tubular epithelial damage

Primary renal proximal tubule epithelial cells (PCS-400-010, ATCC) in a 24-well plate were loaded with ^51^Cr overnight. The next day, the cells were incubated with 10 μM Fer-1 or diluent, and then infected with *C. albicans* at a multiplicity of infection of 10. After 8 hours, the medium above the cells was collected and the epithelial cells were lysed with RadiacWash (Biodex). The amount of ^51^Cr released into the medium and remaining in the cells was determined with a gamma counter, and the percentage of ^51^Cr released in the infected cells we compared to the release by uninfected epithelial cells. The experiment was performed 3 times in triplicate.

### Cytokine measurement *in vitro*

BMDMs were stimulated for 6 and 12 hours with C. albicans (MOI 1). RTECs were simulated with C. albicans for 12 hours (MOI 5). BMDCs were stimulated with curdlan (50 μg/mL; Invivogen) or LPS (1 μg/mL, Sigma Aldrich) for 24 hours. Supernatant were collected and cytokines were measured with Luminex Bead array (R&D Systems) or ELISA (TNFα #DY410, and IL-23 #D2300B; R&D Systems). To determine the effect of JAK and PPAR signaling in DCs, BMDCs were incubated with Ruxolitinib (1µM; Selleck Chemicals), Rosiglitazone (10 μM; Selleck Chemicals), and GW9662 (10 μM; Selleck Chemicals) for 1 hour prior stimulation.

### Quantification and statistical analysis

At least three biological replicates were performed for all *in vitro* experiments unless otherwise indicated. Data were compared by Mann-Whitney corrected for multiple comparisons using GraphPad Prism (v. 9) software. P values < 0.05 were considered statistically significant.

## Data Availability

The authors declare that the data supporting the findings of this study are available within the paper, the accompanying supplementary information files, and the source data (Source Data file).

## Acknowledgements

MS is supported by NIH grant R00DE026856, an American Association of Immunologists Careers in Immunology Fellowship, and a Lundquist Seed grant. MSL is supported by the Division of Intramural Research of the NIAID. RTW is supported by NIH grant R15AI133415, and NJ by the Francis family foundation and by NIH National Center for Advancing Translational Science (NCATS) UCLA CTSI Grant Number KL2TR001882. The content is solely the responsibility of the authors and does not necessarily represent the official views of the National Institutes of Health.

We thank members of the Division of Infectious Diseases at Harbor-UCLA Medical Center for critical suggestions.

## Author contributions

NM and MS designed the experiments. NM, NVS, DA, NJ, and MS performed the experiments. NM, NVS, DA, NJ, and MS analyzed the data. MSL and RTW provided methodology. MS wrote the manuscript. All authors reviewed and edited.

## Notes

### Competing Interest Statement

The authors have declared no competing interest.

## Literature

1. Netea, M.G., Joosten, L.A., van der Meer, J.W., Kullberg, B.J. & van de Veerdonk, F.L. Immune defence against Candida fungal infections. Nat Rev Immunol 15, 630–42 (2015).

2. Lionakis, M.S., Iliev, I.D. & Hohl, T.M. Immunity against fungi. JCI Insight 2, 93156 (2017).

3. Jawale, C.V. & Biswas, P.S. Local antifungal immunity in the kidney in disseminated candidiasis. Current Opinion in Microbiology 62, 1–7 (2021).

4. Brown, G.D. Innate antifungal immunity: the key role of phagocytes. Annu Rev Immunol 29, 1–21 (2011).

5. Dambuza, I.M., Levitz, S.M., Netea, M.G. & Brown, G.D. Fungal Recognition and Host Defense Mechanisms. Microbiol Spectr 5, 0050–2016 (2017).

6. Branzk, N. et al. Neutrophils sense microbe size and selectively release neutrophil extracellular traps in response to large pathogens. Nat Immunol 15, 1017–25 (2014).

7. Drummond, R.A. et al. CARD9-Dependent Neutrophil Recruitment Protects against Fungal Invasion of the Central Nervous System. PLoS Pathog 11, e1005293 (2015).

8. Whitney, P.G. et al. Syk signaling in dendritic cells orchestrates innate resistance to systemic fungal infection. PLoS Pathog 10, e1004276 (2014).

9. Jawale, C.V. et al. Restoring glucose uptake rescues neutrophil dysfunction and protects against systemic fungal infection in mouse models of kidney disease. Sci Transl Med 12(2020).

10. Conti, H.R. et al. Th17 cells and IL-17 receptor signaling are essential for mucosal host defense against oral candidiasis. J Exp Med 206, 299–311 (2009).

11. Conti, H.R. et al. IL-17 Receptor Signaling in Oral Epithelial Cells Is Critical for Protection against Oropharyngeal Candidiasis. Cell Host Microbe 20, 606–617 (2016).

12. Swidergall, M. & Filler, S.G. Oropharyngeal Candidiasis: Fungal Invasion and Epithelial Cell Responses. PLoS Pathog 13, e1006056 (2017).

13. Verma, A., Gaffen, S.L. & Swidergall, M. Innate Immunity to Mucosal Candida Infections. J Fungi (Basel) 3, 60 (2017).

14. Puel, A. et al. Inborn errors of human IL-17 immunity underlie chronic mucocutaneous candidiasis. Curr Opin Allergy Clin Immunol 12, 616–22 (2012).

15. Lionakis, M.S. & Levitz, S.M. Host Control of Fungal Infections: Lessons from Basic Studies and Human Cohorts. Annu Rev Immunol 36, 157–191 (2018).

16. Lionakis, M.S., Netea, M.G. & Holland, S.M. Mendelian genetics of human susceptibility to fungal infection. Cold Spring Harb Perspect Med 4(2014).

17. Swidergall, M. et al. Candidalysin Is Required for Neutrophil Recruitment and Virulence During Systemic Candida albicans Infection. J Infect Dis 220, 1477–1488 (2019).

18. Carpino, N., Naseem, S., Frank, D.M. & Konopka, J.B. Modulating Host Signaling Pathways to Promote Resistance to Infection by Candida albicans. Frontiers in Cellular and Infection Microbiology 7(2017).

19. Lionakis, M.S. et al. Chemokine receptor Ccr1 drives neutrophil-mediated kidney immunopathology and mortality in invasive candidiasis. PLoS Pathog 8, e1002865 (2012).

20. Pappas, P.G., Lionakis, M.S., Arendrup, M.C., Ostrosky-Zeichner, L. & Kullberg, B.J. Invasive candidiasis. Nat Rev Dis Primers 4, 18026 (2018).

21. Lone, S.A., Wani, M.Y., Fru, P. & Ahmad, A. Cellular apoptosis and necrosis as therapeutic targets for novel Eugenol Tosylate Congeners against Candida albicans. Scientific Reports 10, 1191 (2020).

22. Bertheloot, D., Latz, E. & Franklin, B.S. Necroptosis, pyroptosis and apoptosis: an intricate game of cell death. Cellular & Molecular Immunology 18, 1106–1121 (2021).

23. Jorgensen, I., Rayamajhi, M. & Miao, E.A. Programmed cell death as a defence against infection. Nature reviews. Immunology 17, 151–164 (2017).

24. Linkermann, A. et al. Regulated cell death in AKI. J Am Soc Nephrol 25, 2689–701 (2014).

25. Tang, D., Kang, R., Berghe, T.V., Vandenabeele, P. & Kroemer, G. The molecular machinery of regulated cell death. Cell Research 29, 347–364 (2019).

26. Kim, E.H., Wong, S.W. & Martinez, J. Programmed Necrosis and Disease:We interrupt your regular programming to bring you necroinflammation. Cell Death Differ 26, 25–40 (2019).

27. Dhuriya, Y.K. & Sharma, D. Necroptosis: a regulated inflammatory mode of cell death. Journal of Neuroinflammation 15, 199 (2018).

28. Chen, X., Kang, R., Kroemer, G. & Tang, D. Ferroptosis in infection, inflammation, and immunity. J Exp Med 218(2021).

29. Miao, E.A. et al. Caspase-1-induced pyroptosis is an innate immune effector mechanism against intracellular bacteria. Nature immunology 11, 1136–1142 (2010).

30. Wellington, M., Koselny, K., Sutterwala, F.S. & Krysan, D.J. Candida albicans triggers NLRP3-mediated pyroptosis in macrophages. Eukaryot Cell 13, 329–40 (2014).

31. Li, T. et al. TSC1 Suppresses Macrophage Necroptosis for the Control of Infection by Fungal Pathogen <em>Candida albicans</em>. ImmunoHorizons 5, 90–101 (2021).

32. Gross, O. et al. Syk kinase signalling couples to the Nlrp3 inflammasome for anti-fungal host defence. Nature 459, 433–436 (2009).

33. Swidergall, M., Solis, N.V., Lionakis, M.S. & Filler, S.G. EphA2 is an epithelial cell pattern recognition receptor for fungal beta-glucans. Nat Microbiol 3, 53–61 (2018).

34. Swidergall, M. et al. EphA2 Is a Neutrophil Receptor for Candida albicans that Stimulates Antifungal Activity during Oropharyngeal Infection. Cell Rep 28, 423–433 e5 (2019).

35. Sun, W. et al. Cutting Edge: EPHB2 Is a Coreceptor for Fungal Recognition and Phosphorylation of Syk in the Dectin-1 Signaling Pathway. J Immunol 206, 1419–1423 (2021).

36. Chen, J. et al. TAGAP instructs Th17 differentiation by bridging Dectin activation to EPHB2 signaling in innate antifungal response. Nature Communications 11, 1913 (2020).

37. Aaron, P.A., Jamklang, M., Uhrig, J.P. & Gelli, A. The blood-brain barrier internalises Cryptococcus neoformans via the EphA2-tyrosine kinase receptor. Cell Microbiol 20(2018).

38. Kottom, T.J., Schaefbauer, K., Carmona, E.M. & Limper, A.H. EphA2 is a Lung Epithelial Cell Receptor for Pneumocystis β-glucans. J Infect Dis (2021).

39. Swidergall, M. Candida albicans at Host Barrier Sites: Pattern Recognition Receptors and Beyond. Pathogens 8, 40 (2019).

40. Swidergall, M. et al. Activation of EphA2-EGFR signaling in oral epithelial cells by Candida albicans virulence factors. PLOS Pathogens 17, e1009221 (2021).

41. Phan, Q.T. et al. The Globular C1q Receptor Is Required for Epidermal Growth Factor Receptor Signaling during Candida albicans Infection. mBio 12, e0271621 (2021).

42. Höft, M.A., Hoving, J.C. & Brown, G.D. Signaling C-Type Lectin Receptors in Antifungal Immunity. Curr Top Microbiol Immunol 429, 63–101 (2020).

43. Brown, G.D. & Gordon, S. Immune recognition. A new receptor for beta-glucans. Nature 413, 36–7 (2001).

44. Brown, G.D. Dectin-1: a signalling non-TLR pattern-recognition receptor. Nat Rev Immunol 6, 33–43 (2006).

45. de Saint-Vis, B. et al. Human dendritic cells express neuronal Eph receptor tyrosine kinases: role of EphA2 in regulating adhesion to fibronectin. Blood 102, 4431–40 (2003).

46. Finney, A.C. et al. EphA2 Expression Regulates Inflammation and Fibroproliferative Remodeling in Atherosclerosis. Circulation 136, 566–582 (2017).

47. Navarathna, D.H., Roberts, D.D., Munasinghe, J. & Lizak, M.J. Imaging Candida Infections in the Host. Methods Mol Biol 1356, 69–78 (2016).

48. Dunker, C. et al. Rapid proliferation due to better metabolic adaptation results in full virulence of a filament-deficient Candida albicans strain. Nat Commun 12, 3899 (2021).

49. Lionakis, M.S., Lim, J.K., Lee, C.C. & Murphy, P.M. Organ-specific innate immune responses in a mouse model of invasive candidiasis. J Innate Immun 3, 180–99 (2011).

50. Norice, C.T., Smith, F.J., Jr., Solis, N., Filler, S.G. & Mitchell, A.P. Requirement for Candida albicans Sun41 in biofilm formation and virulence. Eukaryot Cell 6, 2046–55 (2007).

51. Liu, Y., Mittal, R., Solis, N.V., Prasadarao, N.V. & Filler, S.G. Mechanisms of Candida albicans trafficking to the brain. PLoS Pathog 7, e1002305 (2011).

52. Carvalho, A. et al. Immunity and Tolerance to Fungi in Hematopoietic Transplantation: Principles and Perspectives. Frontiers in Immunology 3(2012).

53. Leavy, O. Macrophages: Early antifungal defence in kidneys. Nat Rev Immunol 14, 6–7 (2014).

54. Jae-Chen, S. et al. Mechanism underlying renal failure caused by pathogenic Candida albicans infection. Biomed Rep 3, 179–182 (2015).

55. Spellberg, B., Ibrahim, A.S., Edwards, J.E., Jr. & Filler, S.G. Mice with disseminated candidiasis die of progressive sepsis. J Infect Dis 192, 336–43 (2005).

56. Singer, E. et al. Neutrophil gelatinase-associated lipocalin: pathophysiology and clinical applications. Acta Physiol (Oxf) 207, 663–72 (2013).

57. Duggan, S., Leonhardt, I., Hunniger, K. & Kurzai, O. Host response to Candida albicans bloodstream infection and sepsis. Virulence 6, 316–26 (2015).

58. Su, L., Liu, D., Chai, W., Liu, D. & Long, Y. Role of sTREM-1 in predicting mortality of infection: a systematic review and meta-analysis. BMJ Open 6, e010314 (2016).

59. Jang, H.R. & Rabb, H. Immune cells in experimental acute kidney injury. Nature Reviews Nephrology 11, 88–101 (2015).

60. Dimitrova, P., Gyurkovska, V., Shalova, I., Saso, L. & Ivanovska, N. Inhibition of zymosan-induced kidney dysfunction by tyrphostin AG-490. Journal of inflammation (London, England) 6, 13–13 (2009).

61. Khounlotham, M., Subbian, S., Smith, R., 3rd, Cirillo, S.L. & Cirillo, J.D. Mycobacterium tuberculosis interferes with the response to infection by inducing the host EphA2 receptor. J Infect Dis 199, 1797–806 (2009).

62. Zhang, Y. et al. mTORC1 couples cyst(e)ine availability with GPX4 protein synthesis and ferroptosis regulation. Nature Communications 12, 1589 (2021).

63. Dixon, S.J. & Stockwell, B.R. The Hallmarks of Ferroptosis. Annual Review of Cancer Biology 3, 35–54 (2019).

64. Yang, W.S. & Stockwell, B.R. Ferroptosis: Death by Lipid Peroxidation. Trends in cell biology 26, 165–176 (2016).

65. Dixon, S.J. et al. Ferroptosis: an iron-dependent form of nonapoptotic cell death. Cell 149, 1060–72 (2012).

66. Horwath, M.C. et al. Antifungal Activity of the Lipophilic Antioxidant Ferrostatin-1. Chembiochem 18, 2069–2078 (2017).

67. Nur, S. et al. IL-23 supports host defense against systemic Candida albicans infection by ensuring myeloid cell survival. PLoS Pathog 15, e1008115 (2019).

68. Liu, Q. & Wang, K. The induction of ferroptosis by impairing STAT3/Nrf2/GPx4 signaling enhances the sensitivity of osteosarcoma cells to cisplatin. Cell Biol Int 43, 1245–1256 (2019).

69. Brown, C.W., Amante, J.J., Goel, H.L. & Mercurio, A.M. The α6β4 integrin promotes resistance to ferroptosis. The Journal of cell biology 216, 4287–4297 (2017).

70. Legrand, A.J., Konstantinou, M., Goode, E.F. & Meier, P. The Diversification of Cell Death and Immunity: Memento Mori. Mol Cell 76, 232–242 (2019).

71. Li, J. et al. Ferroptosis: past, present and future. Cell Death & Disease 11, 88 (2020).

72. Tang, D., Chen, X., Kang, R. & Kroemer, G. Ferroptosis: molecular mechanisms and health implications. Cell Research 31, 107–125 (2021).

73. Medzhitov, R., Schneider, D.S. & Soares, M.P. Disease tolerance as a defense strategy. Science 335, 936–41 (2012).

74. Doll, S. & Conrad, M. Iron and ferroptosis: A still ill-defined liaison. IUBMB Life 69, 423–434 (2017).

75. Gaschler, M.M. & Stockwell, B.R. Lipid peroxidation in cell death. Biochemical and biophysical research communications 482, 419–425 (2017).

76. Harry, R.S. et al. Metabolic impact of 4-hydroxynonenal on macrophage-like RAW 264.7 function and activation. Chem Res Toxicol 25, 1643–51 (2012).

77. Strasser, D. et al. Syk kinase-coupled C-type lectin receptors engage protein kinase C-δ to elicit Card9 adaptor-mediated innate immunity. Immunity 36, 32–42 (2012).

78. Kapralov, A.A. et al. Redox lipid reprogramming commands susceptibility of macrophages and microglia to ferroptotic death. Nature Chemical Biology 16, 278–290 (2020).

79. Sun, T. & Chi, J.T. Regulation of ferroptosis in cancer cells by YAP/TAZ and Hippo pathways: The therapeutic implications. Genes Dis 8, 241–249 (2021).

80. Jaumouillé, V. & Waterman, C.M. Physical Constraints and Forces Involved in Phagocytosis. Frontiers in immunology 11, 1097–1097 (2020).

81. Bain, J.M. et al. Immune cells fold and damage fungal hyphae. Proceedings of the National Academy of Sciences 118, e2020484118 (2021).

82. Cheng, S.-C. et al. mTOR-and HIF-1α-mediated aerobic glycolysis as metabolic basis for trained immunity. Science (New York, N.Y.) 345, 1250684–1250684 (2014).

83. Vucetic, M., Daher, B., Cassim, S., Meira, W. & Pouyssegur, J. Together we stand, apart we fall: how cell-to-cell contact/interplay provides resistance to ferroptosis. Cell Death & Disease 11, 789 (2020).

84. Khader, S.A. & Thirunavukkarasu, S. The Tale of IL-12 and IL-23: A Paradigm Shift. The Journal of Immunology 202, 629–630 (2019).

85. Thompson, A. & Orr, S.J. Emerging IL-12 family cytokines in the fight against fungal infections. Cytokine 111, 398–407 (2018).

86. Kim, H.S. et al. Curdlan activates dendritic cells through dectin-1 and toll-like receptor 4 signaling. Int Immunopharmacol 39, 71–78 (2016).

87. Parham, C. et al. A receptor for the heterodimeric cytokine IL-23 is composed of IL-12Rbeta1 and a novel cytokine receptor subunit, IL-23R. J Immunol 168, 5699–708 (2002).

88. Oppmann, B. et al. Novel p19 protein engages IL-12p40 to form a cytokine, IL-23, with biological activities similar as well as distinct from IL-12. Immunity 13, 715–25 (2000).

89. Chen, Y. et al. A study on the risk of fungal infection with tofacitinib (CP-690550), a novel oral agent for rheumatoid arthritis. Scientific Reports 7, 6779 (2017).

90. Tsirigotis, P. et al. Treatment of Experimental Candida Sepsis with a Janus Kinase Inhibitor Controls Inflammation and Prolongs Survival. Antimicrobial agents and chemotherapy 59, 7367–7373 (2015).

91. Yang, X.O. et al. STAT3 regulates cytokine-mediated generation of inflammatory helper T cells. J Biol Chem 282, 9358–9363 (2007).

92. Lee, P.W. et al. IL-23R-activated STAT3/STAT4 is essential for Th1/Th17-mediated CNS autoimmunity. JCI insight 2, e91663 (2017).

93. Yao, R. et al. Pathogenic effects of inhibition of mTORC1/STAT3 axis facilitates Staphylococcus aureus-induced pyroptosis in human macrophages. Cell Communication and Signaling 18, 187 (2020).

94. Smith, A.D. et al. Autocrine IL6-Mediated Activation of the STAT3-DNMT Axis Silences the TNFα-RIP1 Necroptosis Pathway to Sustain Survival and Accumulation of Myeloid-Derived Suppressor Cells. Cancer Res 80, 3145–3156 (2020).

95. Wang, W. et al. CD8+ T cells regulate tumour ferroptosis during cancer immunotherapy. Nature 569, 270–274 (2019).

96. Fadok, V.A. et al. Exposure of phosphatidylserine on the surface of apoptotic lymphocytes triggers specific recognition and removal by macrophages. J Immunol 148, 2207–16 (1992).

97. Klöditz, K. & Fadeel, B. Three cell deaths and a funeral: macrophage clearance of cells undergoing distinct modes of cell death. Cell Death Discovery 5, 65 (2019).

98. Dennehy, K.M., Willment, J.A., Williams, D.L. & Brown, G.D. Reciprocal regulation of IL-23 and IL-12 following co-activation of Dectin-1 and TLR signaling pathways. Eur J Immunol 39, 1379–86 (2009).

99. Wang, H. et al. Targeting EphA2 suppresses hepatocellular carcinoma initiation and progression by dual inhibition of JAK1/STAT3 and AKT signaling. Cell Rep 34, 108765 (2021).

100. Melillo, J.A. et al. Dendritic Cell (DC)-Specific Targeting Reveals Stat3 as a Negative Regulator of DC Function. The Journal of Immunology 184, 2638–2645 (2010).

101. Lionakis, M.S. et al. CX3CR1-dependent renal macrophage survival promotes Candida control and host survival. J Clin Invest 123, 5035–51 (2013).

102. Millet, N., Solis, N.V. & Swidergall, M. Mucosal IgA Prevents Commensal Candida albicans Dysbiosis in the Oral Cavity. Front Immunol 11, 555363 (2020).

